# Host transcriptional response to SARS-CoV-2 infection in COVID-19 patients

**DOI:** 10.1101/2021.05.13.443721

**Authors:** Nitesh Kumar Singh, Surabhi Srivastava, Lamuk Zaveri, Thrilok Chander Bingi, Rajarao Mesipogu, Santosh Kumar, Namami Gaur, Nikhil Hajirnis, Pratheusa Maccha, Sakshi Shambhavi, Shagufta Khan, Mamilla Soujanya, Tulasi Nagabandi, Rakesh K. Mishra, Karthik Bharadwaj Tallapaka, Divya Tej Sowpati

## Abstract

**Background:** One of the most perplexing aspects of infection with the SARS-CoV-2 virus has been the variable response elicited in its human hosts. Investigating the transcriptional changes in individuals affected by COVID-19 can help understand and predict the degree of illness and guide clinical outcomes in diverse backgrounds.

**Methods:** Analysis of host transcriptome variations via RNA sequencing from naso/oropharyngeal swabs of COVID-19 patients.

**Results:** We report strong upregulation of the innate immune response, especially type I interferon pathway, upon SARS-CoV-2 infection. Upregulated genes were subjected to a comparative meta-analysis using global datasets to identify a common network of interferon stimulated and viral response genes that mediate the host response and resolution of infection. A large proportion of mis-regulated genes showed a reduction in expression level, suggesting an overall decrease in host mRNA production. Significantly downregulated genes included those encoding olfactory, taste and neuro-sensory receptors. Many pro-inflammatory markers and cytokines were also downregulated or remained unchanged in the COVID-19 patients. Finally, a large number of non-coding RNAs were identified as down-regulated, with a few of the lncRNAs associated with functional roles in directing the response to viral infection.

**Conclusions:** SARS-CoV-2 infection results in the robust activation of the body’s innate immunity. Reduction of gene expression is well correlated with the clinical manifestations and symptoms of COVID-19 such as the loss of smell and taste, and myocardial and neurological complications. This study provides a critical dataset of genes that will enhance our understanding of the nature and prognosis of COVID-19.

## Introduction

SARS-CoV-2, the coronavirus causing COVID-19 has brought enormous economical, health and human cost to the world. SARS-CoV-2, like MERS-CoV and SARS-CoV that caused epidemics in 2003 and 2012, respectively, is the latest coronavirus to infect humans and can cause severe disease with long term and/or fatal consequences in some individuals [1]. COVID-19 progression is unpredictable and ranges from asymptomatic, mild infection to life-threatening acute respiratory distress syndrome (ARDS) [2]. Mild infections are characterized by symptoms such as fever and cough, and are usually resolved spontaneously in a majority of the infected individuals. However, in severe cases, patients develop ARDS and acute lung injury with damage to alveolar lumen leading to inflammation and pneumonia, and resulting in Intensive Care Unit (ICU) admissions or even death [3]. The cause of such differential outcomes is unknown though older age, presence of other disease comorbidities and the host immune response to the virus are believed to be some of the key factors in disease prognosis.

During a typical infection, the immune system activates two distinct antiviral responses in response to pathogen entry. The first defense relies on the body’s innate immunity and is mediated by transcriptional induction of type I and III interferon (IFN) and the subsequent activation of IFN-stimulated genes (ISGs) [4]. This then triggers the recruitment and coordination of adaptive immune cells via the secretion of chemokines. In some individuals this process can be associated with an excessive increase in pro-inflammatory cytokines. Finally, sometimes the virus may evade the body’s innate immunity, leading to an inadequate or delayed response allowing unrestrained viral replication subsequently leading to hyper inflammatory responses [5].

Recent transcriptomic studies investigating the host response using RNA-sequencing (RNA-seq) have tried to identify a molecular signature of the response to SARS-CoV-2 infection. Single cell RNA-seq performed on human cell lines infected with SARS-CoV-2 has demonstrated that canonical ISGs are broadly induced [6]. The failure to launch an appropriate degree of interferon-mediated response despite virus replication has been shown to be associated with high levels of chemokine induction in some individuals [7]. The reasons underlying the diversity in the immune response and the correlates for severe infection and death due to COVID-19 are still unclear and need to be investigated across diverse ethnicities, age groups and susceptibilities.

In this work, we have identified the changes induced in the transcriptome of individuals positive for COVID-19 infection. Our analysis of samples from Indian patients reveals a massive downregulation of host genes in response to SARS-CoV-2 infection. The up-regulated genes comprised only ~27% of the mis-expressed genes, most of which were part of a strong immune response and innate immunity network. Down-regulated genes were enriched in pathways that are associated with systemic symptoms of COVID-19 such as cardiac, pancreatic and muscle related issues as well as neuronal and sensory abnormalities. We also observed significant down-regulation of smell and taste receptor genes, which correlates well with olfactory dysfunction, now acknowledged as a key symptom of COVID-19. Interestingly, many long noncoding RNA (lncRNAs) were also found mis-regulated, with functional roles in a myriad of functions including mounting a strong antiviral response [8].

## Results and discussion

### Extensive downregulation of host transcriptome in COVID-19 patients

We analyzed total RNA isolated from nasopharyngeal samples of 36 COVID-19 positive patients from the southern state of Telangana, India, along with 5 COVID-19 negative samples as controls (see Methods and Supplementary table 1). Principal component analysis (Figure 1A) shows that the patient samples did not cluster very well together, suggesting high sample variability and possibly a diverse transcriptomic response by individuals against SARS-CoV-2. However, there was a clear distinction between the positive and the control samples and a total of 56,640 genes were analyzed for differential expression (methods).

**Figure 1 :**
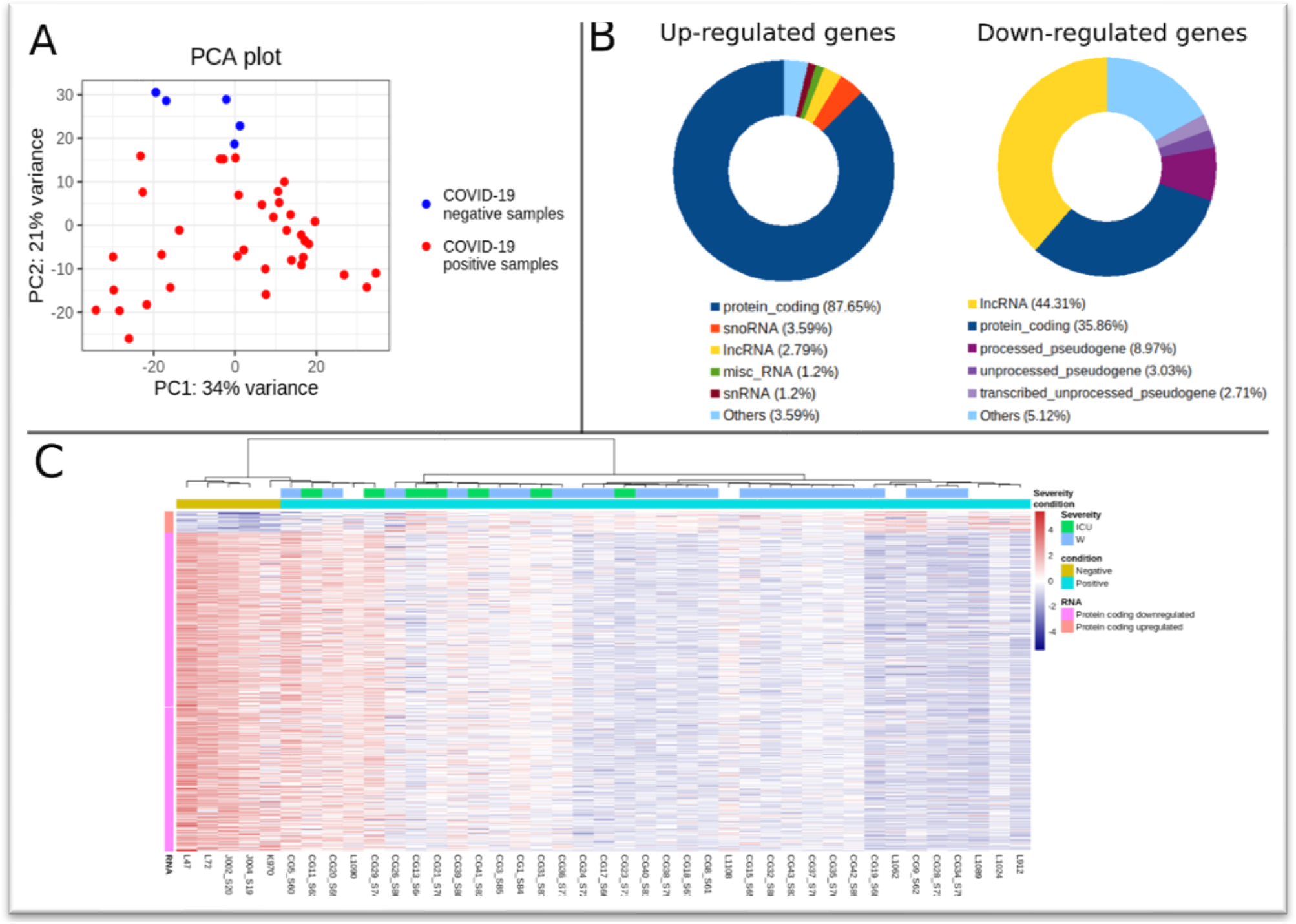
Differentially expressed genes (DEGs) and their expression in COVID-19 patients compared to controls. (A) Principal component analysis representing the 36 COVID-19 positive (red) and 5 negative samples (blue). Principal component 1 plotted on the x-axis explains 34% of the variance and principal component 2 on the y-axis explains 21% variance in the transcriptomic profiles. (B) Pie chart of biotype composition of DEGs. Only the top 5 categories for up-regulated and down-regulated genes are shown. Most of the up-regulated genes are protein-coding but the highest percentage of down-regulated genes are lncRNAs. (C) Heatmap of the significantly expressed protein coding genes (rlog transformed expression values). Each bar marks the expression level of a gene from highest (red) to lowest (blue) as per the scale on the right. Sample names are indicated on the x-axis (clustered by negative and then positive samples) and their metadata is shown on the top. ‘Condition’ indicates COVID-19 negative (gold bar) or positive (blue bar) status while ‘Severity’ indicates whether the patients needed ICU intervention (ICU, green bars) or were discharged from the general ward (W, dark blue bars). The Y-axis bar on the left marks the number of up-regulated protein coding genes (orange) followed by the down-regulated protein coding genes (pink) in patients compared to the controls.

We identified 9319 differentially expressed genes (DEGs), of which only 251 genes were up-regulated while 9068 genes were down-regulated in COVID-19 patients (Supplementary table 2). While most of the up-regulated DEGs were protein coding (220 genes), only 35% (3252 genes) of the down-regulated DEGs were protein coding (Figure 1B and Supplementary table 3). The levels of expression of all the 3472 differentially regulated protein coding genes are summarized across the samples in Figure 1C. A large proportion of the down-regulated DEGs were non-coding RNAs, lncRNAs being the biggest contributors (4018 lncRNAs). The large number of differentially expressed lncRNAs suggests a key role for them during viral infection as described in earlier studies [8, 9].

The downregulation of a large proportion of the DEGs is an interesting observation that may be associated with the phenomenon of viral control and switching off host protein production [10]. Besides affecting the immune response, the host cell proteins fade and viral proteins become more prominent with the progress of the infection. The precise mechanisms remain unclear but host shut-off is likely brought about at multiple levels, including down-regulation of host transcription and at the level of translation, where the mis-regulated lncRNAs may also play a role.

### Up-regulation of immune related genes and the interferon-mediated immune response

We found that protein coding up-regulated genes were enriched mostly for immune system related GO terms and pathways, especially those associated with Type I interferon signaling and defense response to viral infections (Figure 2A). Prominent clusters of genes associated with multiple viral infections could also be identified (Figure 2B), confirming the upregulation of infection response pathways. Systemic effects on multiple organs, a hallmark of COVID-19 and associated with high-risk comorbidities such as diabetes and thyroid dysfunction, are indicated. Viral myocarditis appears to be induced either as a direct result of the infection or as a downstream consequence of the immune response activation. A strong association between myocardial injury and significantly worse clinical course and increased mortality has been documented and there have been several cases of acute and/ or long-term myocarditis following COVID-19 [11, 12]. The complete list of significantly enriched GO terms and KEGG pathways is provided in Supplementary tables 4 and 5 respectively.

**Figure 2 :**
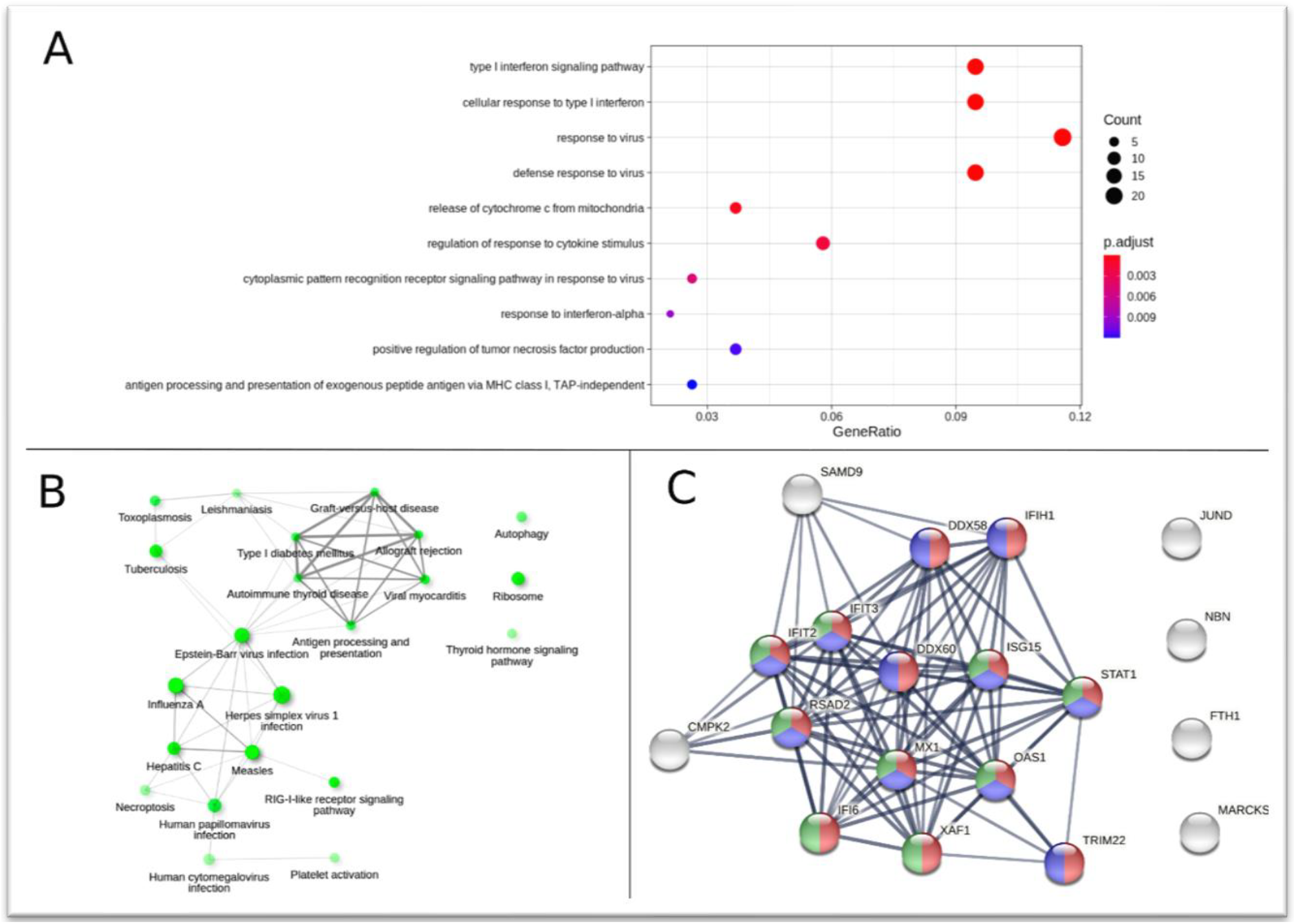
Functional enrichment analysis for protein coding up-regulated genes (220) in COVID-19 patients compared to controls. (A) Top 10 enriched GO terms associated with upregulated genes, based on adjusted p-value (p.adj) values. Size of dots represents the number of genes and color signifies p.adj value. (B) Interaction network showing enriched KEGG pathways. Nodes sharing 30% or more genes are connected by edges whose thickness represents percentage of common genes. Size of the node represents the number of genes in that pathway (ranging from 17 to 4) and shade indicates the significance, darker shades being more significant. (C) Protein-protein interactions of 19 genes showing a tightly connected network involved in “innate immune response” (red nodes), “defense response to virus” (blue nodes) and “type I interferon signalling pathway” (green nodes). These genes were up-regulated during COVID-19 infection in our analysis as well as in other published datasets. Edges depict both functional and physical associations. Edge thickness indicates the confidence in the interaction. All active interaction sources in the STRING database are considered. The minimum interaction score for an edge is set at a high confidence level of 0.7.

We performed a meta-analysis of published datasets (see methods) to identify a set of 19 upregulated genes that showed increased expression in our analysis as well as in other studies on COVID-19 samples (Supplementary Table 6). Protein-protein interaction (PPI) network analysis showed that most of these genes are highly interconnected (Figure 2C), involved in “innate immune response” (red nodes), the “type I interferon signaling pathway” (green nodes), and/or “defense response to virus” (blue nodes). Interferon (IFN)-induced antiviral proteins play a crucial role in distinguishing self and non-self mRNAs during viral infection by inhibiting viral mRNA expression which lacks 2'-O-methylation of the 5’ cap. *IFI6*, recently proposed to be a part of a novel transcriptional signature for COVID-19 infected cells [13], formed one of the key PPI nodes along with *IFIT* family gene paralogs that restrict viral translation and modulate viral response pathways [14]. *IFIH1* and *ISG15* that drive the innate immune response upon sensing viral RNA were also identified as key upregulated genes [15]. The induction of these interferon stimulated genes (ISGs) appears concomitant with the upregulation of *STAT1* in these patients. STAT1 binding to ISGs mediates the interferon triggered host response and its dysfunction has been associated with hyperactivation of inflammatory pathways in individuals with acute COVID-19 pathophysiology [16]. IFN mediated activation of the JAK-STAT signaling pathway also has a role in inducing necroptosis, found enriched in our pathway analysis and associated with ARDS development in some patients [17].

We identified a few other genes involved in viral response that were upregulated in patients with active COVID-19 infection. Increased *OAS1* levels confer protection against severe COVID-19 [18] and this family of antiviral genes has been shown to be associated with the host response [19]. *OAS1* was significantly up-regulated in our study (log2FC = 1.51) although *OAS2* and *OAS3*, part of the same cluster, were only moderately upregulated. The other genes in our network included *XAF1* and *MX1*, upregulated pro-viral factors recently identified in SARS-CoV-2 infection [20, 21]. *MX1/2* and *RSAD2* - another ISG identified in our analysis - play a role in restricting many viral infections along with OAS proteins, although their role in SARS-CoV-2 immunity is not yet defined [22]. Antiviral genes *CMPK2* (IFN induced HIV restriction factor [23]), *TRIM22* (antiviral activity against other RNA viruses [24]), and the oxidative stress response gene *FTH1* involved in iron homeostasis (up-regulated in PBMC of convalescent SARS patients [25]) had increased expression. Finally, we also found the DExD/H-Box Helicase antiviral factors (*DDX58* and *DDX60*), that promote RIG-I like receptor mediated signaling, upregulated in the network.

Interestingly, all the MHC class 1 and some of the MHC class 2 genes (*HLA-A,B,C,E,F,* and *HLA-DQB1, DR-B1, DR-B5*) involved in T-cell mediated cell death and the antibody-mediated adaptive immune response, were also upregulated along with *RFX5*, that binds to MHC-II promoters. However, many of the proinflammatory markers (*TNF, IL6, IL1B, CCL3, CCL4* and *CXCL2*) were not significantly changed, suggesting an absence of hyperinflammation in these patients. Overall, our results suggest the activation of a robust innate immune response in COVID-19 patients.

### Down-regulated protein coding genes are associated with systemic and cellular host processes

Many of the down-regulated protein coding genes in our analysis were associated with signaling and ion transport related GO terms. We found significant enrichment for neurotransmission, heart and muscular contraction related terms (Figure 3A), indicating impairment of these cellular processes in the COVID-19 patients. KEGG pathway analysis (Figure 3B) confirmed the association of the downregulated genes in calcium signaling and the RAS and cAMP signaling pathways, which may help drive viral replication. The RAS signaling pathway may be involved in pathogenesis of COVID-19 and plays an important role in cardiovascular diseases, neurodegenerative diseases, and acute lung injury [26]. We also noticed enrichment of a group of cardiac function related pathways (Figure 3B). Many of the genes from these KEGG pathways were related to cellular calcium signaling (CACNs) and were down-regulated. The key cardiac proteins, troponin and tropomyosin, were also down-regulated. Calcium ions are required for proper functioning of troponin and tropomyosin dependent cardiac muscle contraction. A disruption of cellular calcium signaling together with down-regulated cardiac proteins indicates possibility of myocardial issues in COVID-19 patients. A few of the down-regulated genes were associated with the pancreatic and insulin secretory systems, in agreement with recent work showing that the insulin requirement for patients with diabetes mellitus is considerably higher at the peak of COVID-19 illness [27].

**Figure 3 :**
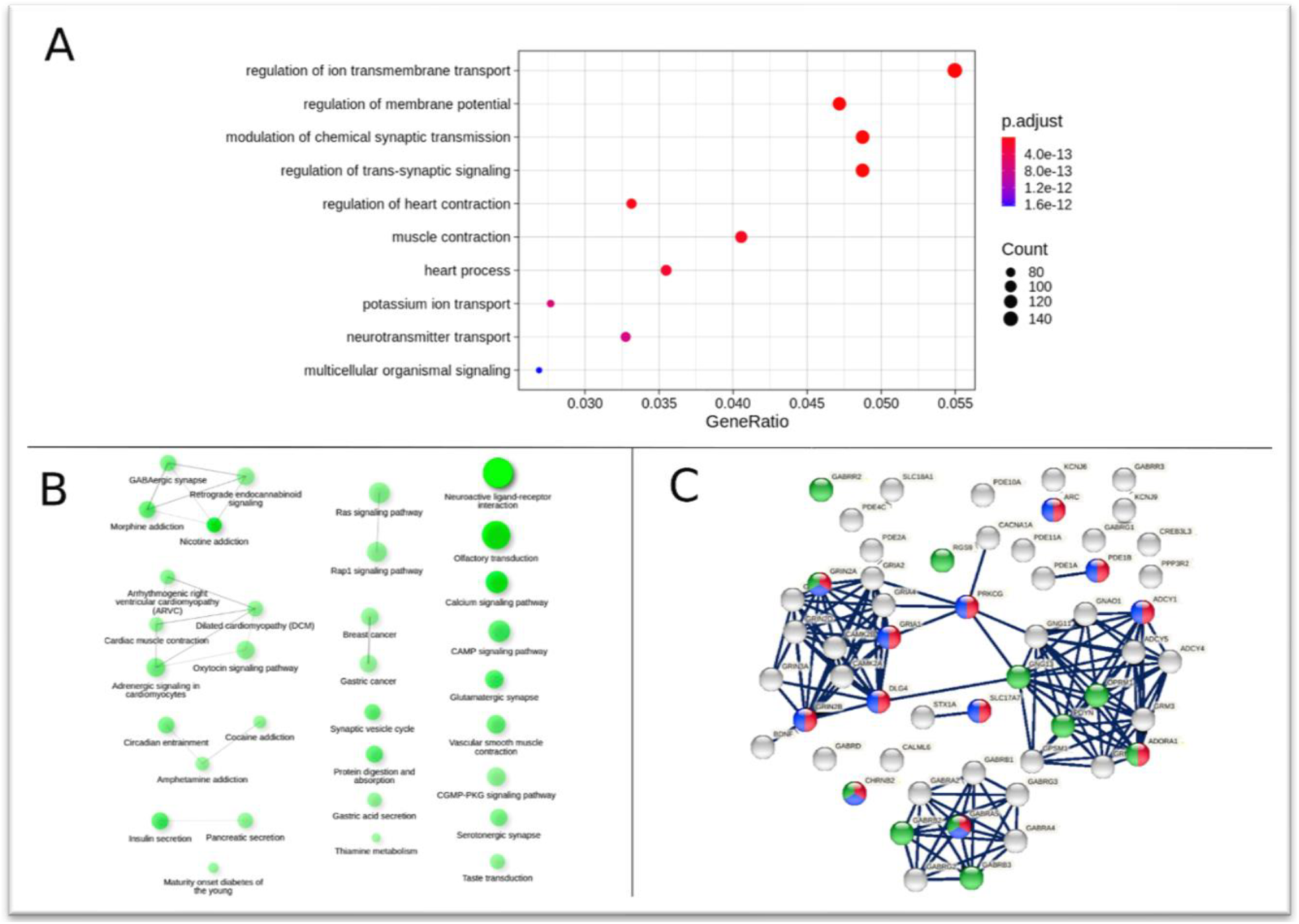
Functional enrichment analysis for protein coding down-regulated genes (3252) in COVID-19 patients compared to controls. (A) Top 10 enriched GO terms associated with downregulated genes, based on adjusted p-value (p.adj) values. Size of dots represents the number of genes and color signifies p.adj value. (B) Interaction network showing enriched KEGG pathways associated with the downregulated genes. Nodes sharing 30% or more genes are connected by edges whose thickness represents percentage of common genes. Size of the node represents the number of genes in that pathway (ranging from 136 to 7) and shade indicates the significance, darker shades being more significant. (C) Protein-protein interactions of 55 genes showing three tightly connected clusters of genes involved in “cognition” (red nodes), “learning and memory” (blue nodes) and “sensory perception” (green nodes). These genes were involved in 4 addiction related pathways. Edges depict both functional and physical associations. Edge thickness indicates the confidence in the interaction. All active interaction sources in the STRING database are considered. The minimum interaction score for an edge is set at a high confidence level of 0.7.

Interestingly, the KEGG pathways “olfactory transduction” and “taste transduction” were found significantly enriched (Figure 3B). 105 out of 118 genes from the “olfactory transduction” pathway, were identified as down-regulated, along with several peripheral sensory receptors, especially the olfactory and taste receptors. With better understanding of the disease over the last year, olfactory dysfunction has emerged as a key symptom of COVID-19 [28] and our results suggest sensory impairment, associated with a loss of smell and taste [29], in these patients. There may also be implications for neuronal infectivity via the olfactory and respiratory tracts. The complete list of enriched GO terms and KEGG pathways associated with the downregulated genes is given in Supplementary tables 7 and 8.

The connection between the downregulation of genes in neurological pathways identified in our analysis and possible neurological associations needs to be further worked out. Cytokines generated in the immune response interacting with various receptors are postulated to be one mechanism to bring about multiple downstream systemic effects [30]. An unexpected finding among genes showing decreased expression in our dataset was the enrichment of drug addiction and neuroactive ligand-receptor pathways (Figure 3B). A PPI analysis (figure 3C) of the 55 genes involved in the four significantly enriched addiction pathways showed a strong network of genes from the family of Gamma-Aminobutyric Acid Type A (GABA) receptors. Reduction in GABA and alterations in GABA receptor levels are associated with stress-induced anxiety and depression phenotypes [31], increasingly recognized in COVID-19 patients. These results are also relevant in the context of the effect of the infection on GABAergic interneurons in the olfactory bulb that connect with the sensory neurons in the olfactory epithelium. Another strongly connected network contained the GRIN genes which are part of the N-methyl-D-aspartate (NMDA) receptors family and are involved in memory, learning and synaptic development. SARS-CoV-2 antibodies generated due to the host immune response can cause neurological symptoms and epilepsy in some patients of COVID-19 due to anti-NMDAR encephalitis [32]. In this context, many of the genes involved in synapse organization and regulation were also downregulated in our analysis highlighting the potential for neurological complications observed in COVID-19 patients [33].

### Systemic downregulation of long non-coding RNAs upon SARS-CoV-2 infection

We identified 4025 lncRNAs mis-regulated in COVID-19, most of them expressed to a lesser extent than the protein coding genes (Supplementary figure S1). A few of these lncRNAs are known to have functional roles during viral infection. For example, *ZBTB11-AS1*, an anti-sense lncRNA to *ZBTB11*, that regulates neutrophil development was up-regulated [34]. Anti-sense lncRNA are associated with regulation of transcriptional gene activation triggered by small RNAs [35], and we found that indeed both *ZBTB11-AS1* and *ZBTB11* were significantly up-regulated in COVID-19 patients (log2FC 2.01 and 1.05, respectively). A few other lncRNAs such as *HEIH* and *IGF2-AS* (associated with recurrence in Hepatitis C virus related hepatocellular carcinoma and Hep C viral replication) [36, 37], *PACER* (positively regulates *COX2* production and *NF-kB* [38]) and *MALAT1* (reduces activation of *NF-kB*, *MAPK* and inflammatory factors) were also significantly mis-regulated.

However, the functional role of many lncRNAs in response against viral infections is not well characterized. We therefore asked if any associated protein-coding genes were affected by the lncRNA mis-regulation. Since a substantial proportion of the functionally annotated lncRNAs are cis-acting [39], we searched for the nearest protein-coding genes to the mis-regulated lncRNAs (methods) and identified 720 differentially expressed protein coding genes. We then grouped the lncRNAs based on distance and strand information in context of the nearest protein-coding genes (Figure 4A). Interestingly, most of the lncRNAs overlap their nearest genes and are on the anti-sense strand. Further, Gene Ontology analysis showed enrichment of terms only in the groups where the lncRNA was located within the gene or in its upstream promoter (Figure 4B). These GO terms were associated mostly with brain or neuron development related processes (Supplementary table 9). Further investigation is needed into the role of the identified lncRNAs in these pathways and their possible association with the complications seen in COVID-19 patients.

**Figure 4 :**
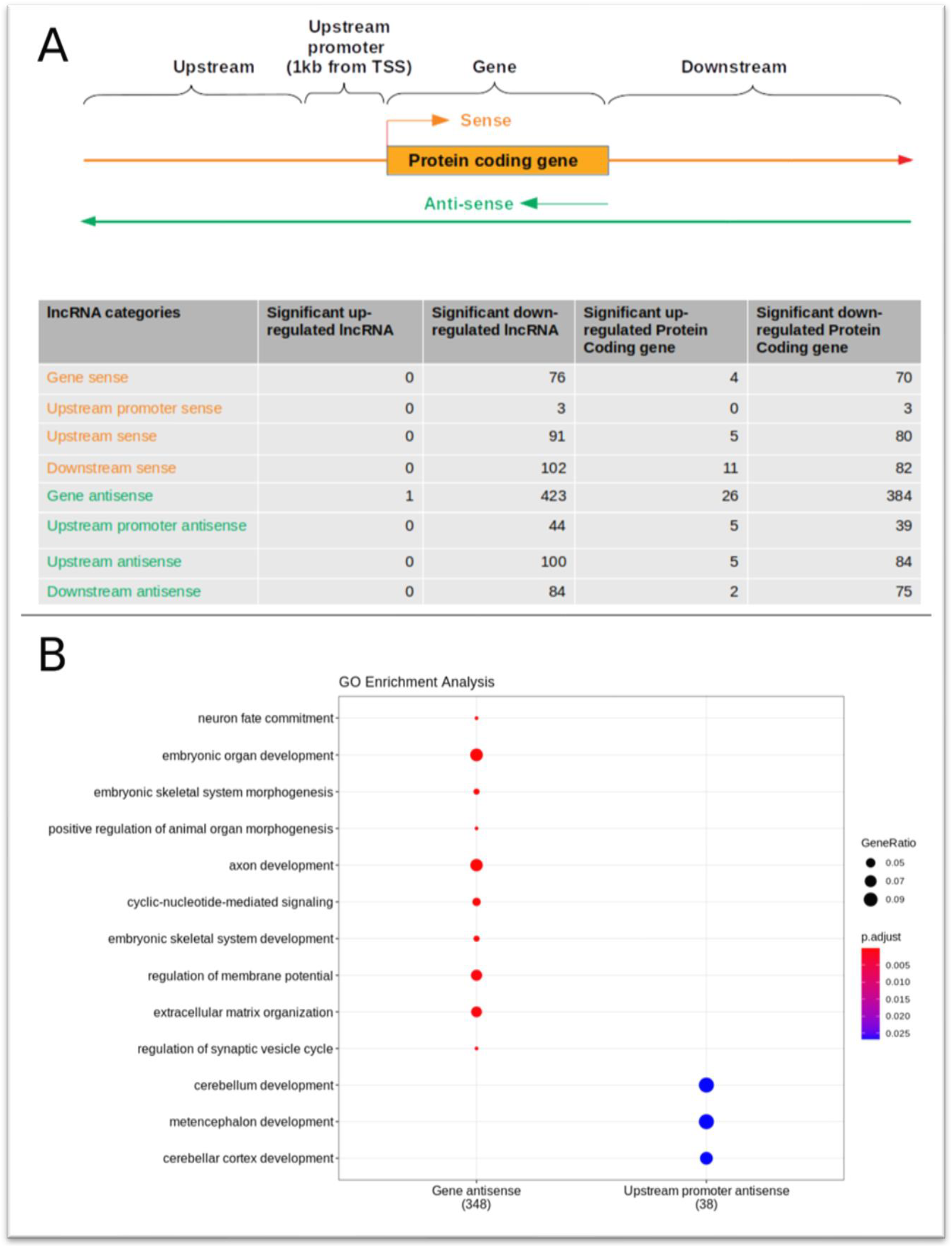
Enrichment analysis of significant protein coding genes closest to lncRNA. (A) Schematic showing the method employed to categorize differentially expressed lncRNAs. The lncRNAs are divided into 8 groups based on their location - genomic strand and distance to their closest differentially expressed protein coding gene. Number of lncRNA and their closest protein coding genes for each category is shown in the table. Most of the lncRNA were on the anti-sense strand and overlapping with genic region or promoter of the closest protein coding gene. (B) GO enrichment analysis of the nearest protein coding genes associated with the differentially expressed lncRNAs. Enriched terms were identified only when the associated lncRNA was present in the genic or promoter contexts, “Gene antisense” and “Upstream promoter antisense”. Only top 10 enriched terms, based on adjusted p-value (p.adj) values, are plotted. Size of dots represents the number of genes and color signifies p.adj value.

## Conclusion

This study identifies the key enriched pathways and processes associated with COVID-19 (summarized in Figure 5). Most of the genes in our analysis were downregulated in the patient samples, suggesting a large-scale systemic response spanning not just lung and respiratory complications but also neurological and cardiac issues, defects in insulin secretion and in the neuro-sensory and olfactory systems. The downregulation of smell and taste pathways points to a tight association of olfactory dysfunction with SARS-CoV-2 infection, even in otherwise asymptomatic individuals.

**Figure 5 :**
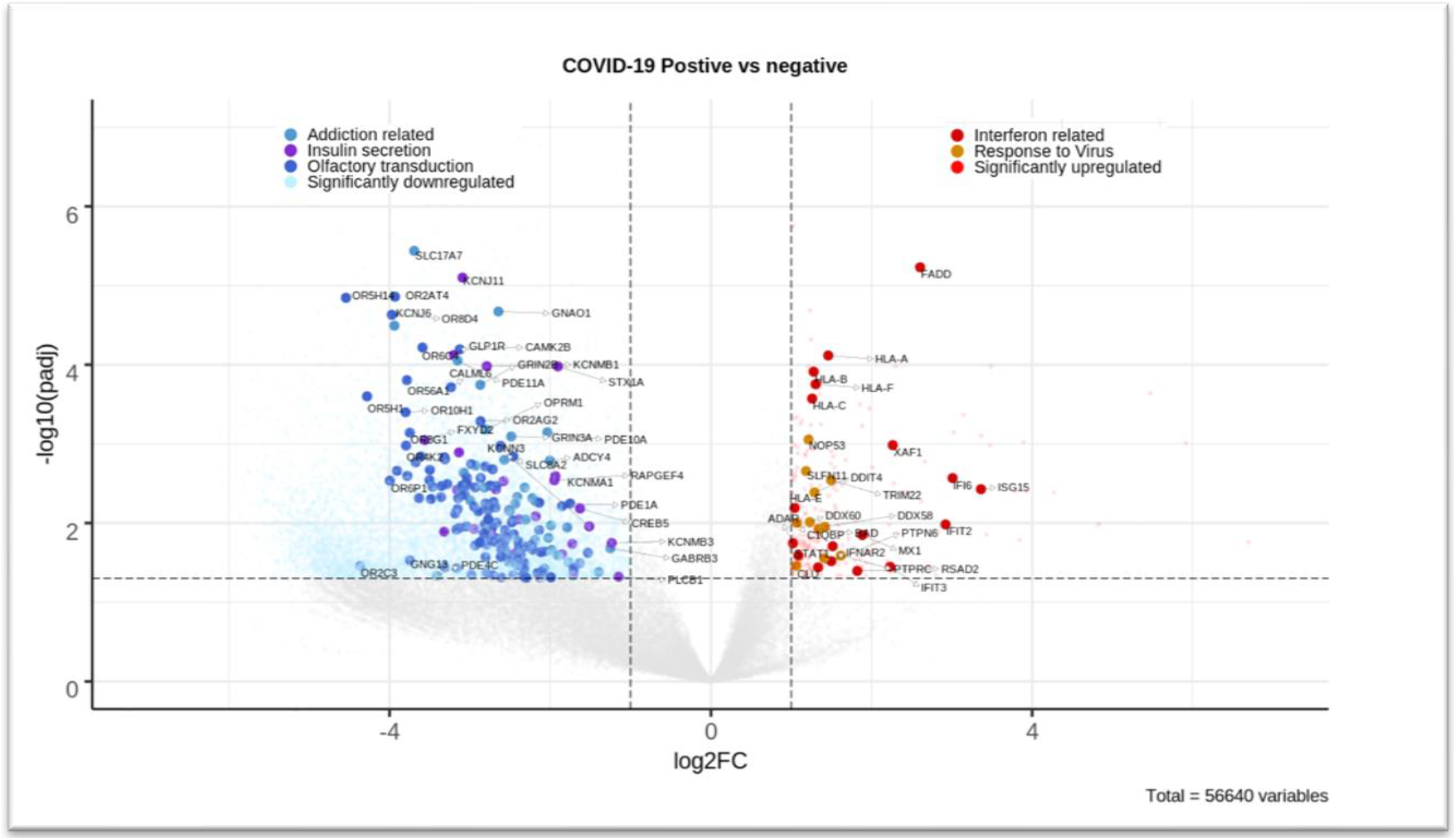
Enriched GO processes and KEGG pathways associated with significant DEGs. Volcano plot showing rlog transformed expression values of all the genes. Each gene is plotted based on the log2 fold change value (X-axis) and mlog10 adjusted p-value (Y-axis). Vertical dashed line (absolute (log2 fold change) = 1) and horizontal dashed line (padj = 0.05) shows the criteria set for defining significant DEGs. Genes not changing significantly are colored grey. Up-regulated genes (colored red) are on the right side of the plot while down-regulated genes (colored blue) are on the left side of the plot. Genes from significantly enriched GO terms and KEGG pathways are highlighted. For example, genes labelled as “Interferon related” are associated with the GO terms “type I interferon signaling pathway” and “cellular response to type I interferon”. Genes labelled as “Response to virus” are associated with the GO terms “defense response to virus” and “response to virus”. Genes marked as “Addiction related” are from 4 enriched KEGG pathways namely “Nicotine addiction”, “Morphine addiction”, “Cocaine addiction” and “Amphetamine addiction”. The other labels correspond to genes from the KEGG pathway “Insulin secretion” and “Olfactory transduction”.

Our results suggest activation of a robust innate immune response, specifically type I interferon pathways, upon SARS-CoV-2 infection. Rapid innate immune activation, usually associated with clearance of the virus, and the absence of pro-inflammatory markers have generally been associated with better disease prognosis. These findings in the context of other health conditions in an individual may help understand varied disease severity. We have identified a significant network of key immune response related genes that are in agreement with several other studies on virus-infected samples, and can provide insights on the immunological perturbations that influence COVID-19 disease progression. Further studies should be in focus to pursue the role of genes and pathways identified from COVID-19 transcriptomic analysis from diverse backgrounds to understand the disease and improve clinical interventions and policies in the pandemic.

## Methods

### Sample metadata

We received 41 samples where 36 samples were confirmed to be COVID-19 positive using RT-PCR method and 5 samples were negative. The age of the patients ranges from 10 to 80 years old. There were 11 females and 30 males in our dataset. We also had severity of COVID-19 for 30 positive patients. 7 patients required intensive care unit (ICU) intervention while 23 were discharged from COVID-19 ward (W). The detailed metadata information about the samples is provided in Supplementary table 1.

### Sample preparation and sequencing

This study was approved by the Institutional Ethics Committee to use patient samples for sequencing and all guidelines were followed. The samples were collected from nasopharyngeal or oropharyngeal swabs as previously described [40]. Total RNA was extracted from 1ml of VTM using TRIzol reagent (Thermo Fisher Scientific, USA) or TRIzol in combination with Direct-zol RNA Microprep Kits (Zymo, USA) according to the manufacturer’s protocol. RNA was quantified using Qubit RNA HS Assay Kit (Thermo Fisher Scientific, USA) and 1 μg total RNA was used for library preparation. Ribosomal RNA was removed using the RiboCop rRNA Depletion Kit (Lexogen, Austria).

RNA-Seq libraries were made using the CORALL Total RNA-Seq Library Prep Kit (Lexogen, Austria) according to the manufacturer’s protocol. Briefly, rRNA depleted RNA was reverse transcribed using displacement stop primers which contain partial Illumina-compatible P7 sequences. Linkers containing partial Illumina-compatible P5 sequences and Unique Molecular Identifiers were ligated to the 3’ end of cDNA fragments. The library was PCR amplified to add the remaining adapter sequences and 12 nucleotide unique indices for multiplexing. Samples were sequenced at PE150 using the Illumina Nova Seq 6000.

### Data Processing and Analysis

Raw sequencing reads were checked for quality using FASTQC v0.11.9 [41] and illumina universal adapters and low-quality reads were removed using cutadapt 2.8 [42]. We removed reads with quality scores less than 20 and discarded reads smaller than 36 bp. The processed reads were then again checked for quality using FASTQC and then mapped to the human genome GRCh38 using STAR 2.7.3a with default parameters [43]. The STAR index was generated using the GRCh38 genome and annotation gtf file downloaded from ENSEMBL [44]. BAM files generated were then sorted according to position using Samtools 1.10 [45]. The quality of BAM was assessed using QualiMap v.2.2.2-dev [46]. All the quality reports were compiled using multiqc 1.9 [47]. Uniquely aligned reads were counted using HTSeq [48]. There were 60683 genes in the gtf file for which we had count information. Genes with read count 0 across all the samples were removed resulting in 56640 genes for further analysis. Differential gene expression analysis was performed using DESeq2_1.24.0 [49]. Genes with adjusted p-value < 0.05 and absolute log2 Fold change > 1 were considered differentially expressed resulting in 9319 genes, remaining 47321 genes were removed from further analysis. For PCA plot and heat map, the raw read counts were rlog normalized, available with the DESeq2 package.

### Functional enrichment analysis

For functional enrichment analysis, clusterProfiler_3.12.0 [50]was used for GO term enrichment and ShinyGO [51] was used for Kyoto Encyclopedia of Genes and Genomes (KEGG) pathway analysis. We only used the Biological process for GO term enrichment analysis. Similar enriched terms were further merged using the ‘simplify’ function of clusterProfiler with similarity cutoff set to 0.7. ‘p.adjust’ was used as a feature to select representative terms and ‘min’ was used to select features. ‘Wang’ was used as a method to measure similarity. For ShinyGO the web tool was run with the default parameter. The complete list of enriched KEGG pathways were downloaded. For better clarity, only the top 20 significant pathways based on FDR were shown in the figures. Edge cut-off was set to 0.3, so edges were created only if two nodes shared at least 30% of the genes. Darker nodes represent more significantly enriched pathways and size of the node represents the number of input genes in that pathway. Thickness of edges was proportional to percentage of overlapping genes.

### Protein-protein interaction network

Protein-protein interaction network was created using STRING-db [52]. Both physical and functional protein associations were considered for edges. All the active interaction sources were considered for generating the network. The medium confidence required for interaction score was set to 0.7. Only input proteins were used for generating the network. Few chosen enriched biological processes or KEGG pathways as reported by STRING-db were shown by color code of the nodes.

### Identifying nearest genes for the lncRNA

We used bedtools (v2.26.0) [53] to find nearest genes for the lncRNA. Bedtools closest searches for overlapping features in two bed files. If there is no overlap, the nearest feature is reported with genomic distance between the two features. Closest features were reported irrespective to strand. The bed file containing differentially expressed lncRNA genomic locations was used as a query bed file (option -a) and a bed file containing all protein coding RNA genomic locations was used as reference bed file (option -b). The lncRNA were classified based on the closest gene. If both the gene and lncRNA were on the same strand, lncRNA was classified as “sense”, otherwise it was classified as “anti-sense”. Next, the lncRNA was classified based on the distance from the closest gene. If the gene and lncRNA were overlapping, lncRNA was classified as “Gene”. If the lncRNA is within 1kb upstream from the closest gene, it is classified as “upstream promoter”. Finally, lncRNA upstream of the closest gene by more than 1 kb was classified as “upstream’ and lncRNA downstream of the gene was classified as “downstream”. This resulted in 8 sets of classification for the lncRNA.

### Meta-analysis of existing transcriptomic data for COVID-19

We performed data mining and compiled a list of 9 published works on transcriptomic host response to COVID-19 [54–62]. We then overlapped our up-regulated/down-regulated genes with a list of genes that were reported to be up-regulated/ down-regulated in the published datasets. For down-regulated genes the overlap was not good between the published dataset. For up-regulated genes, we generated a list of genes that were found to be up-regulated in at least 3 publications. These 19 genes are given in Supplementary table 6.

## Data availability

All the raw RNA-seq data and their respective count data has been submitted to Gene Expression Omnibus (GEO) database and can be accessed under the accession number GSE166530.

## Supplementary data

Supplementary data can be downloaded from the following location https://drive.google.com/file/d/1LhIaewbZ2aZm2QsjElaslh7bAMkuMzR8/view?usp=sharing

---

**Supplementary figure S1:**
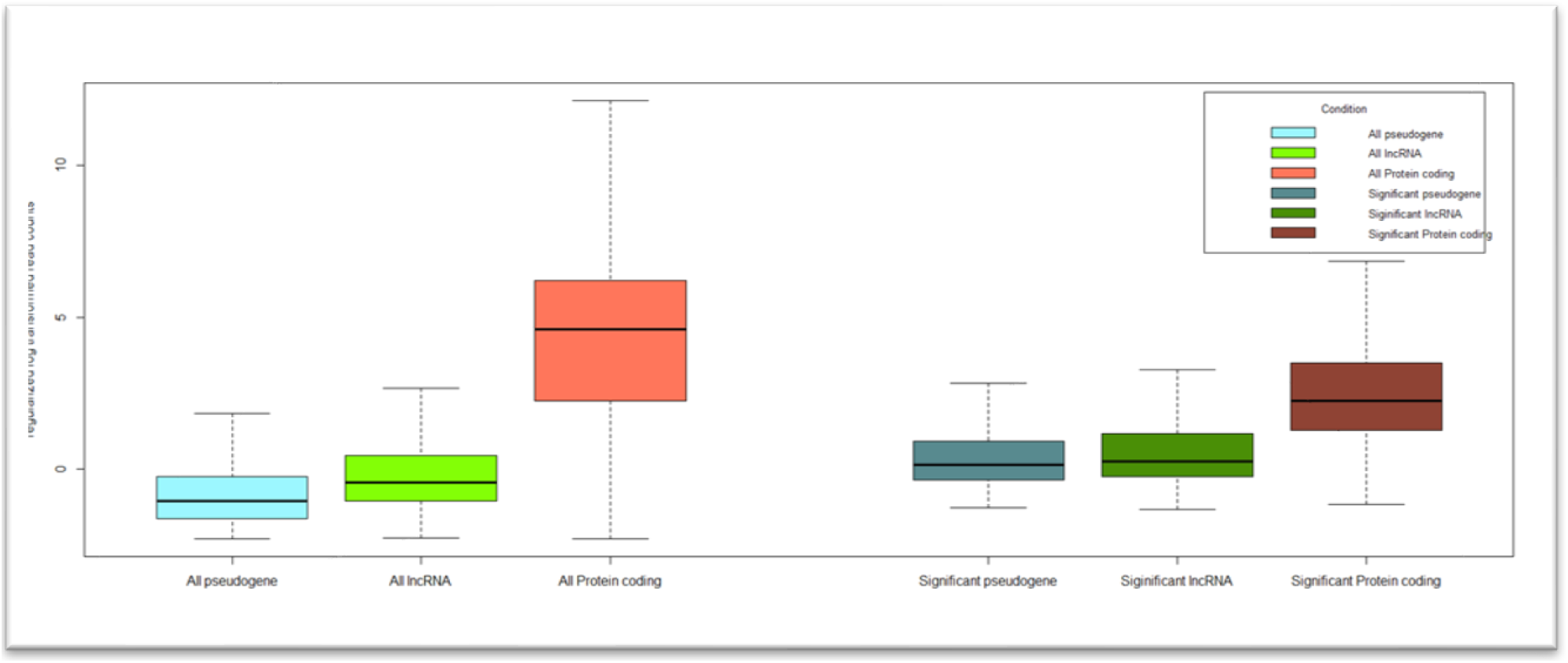
Boxplot showing distribution of expression data for protein-coding genes, lncRNAs and pseudogenes. Distribution of expression value (rlog transformed) for pseudogenes, lncRNAs and protein-coding genes displayed in the form of boxplot. All the pseudogenes, lncRNAs and protein-coding genes are shown on the left side of the plot while differentially expressed are shown on the right side of the plot. Median of the distribution is represented by a horizontal line and whiskers are extended to 1.5 times interquartile range.

